# Particle morphology of medusavirus inside and outside the cells reveals a new maturation process of giant viruses

**DOI:** 10.1101/2021.10.25.465829

**Authors:** Ryoto Watanabe, Chihong Song, Yoko Kayama, Masaharu Takemura, Kazuyoshi Murata

## Abstract

Medusavirus, a giant virus, is phylogenetically closer to eukaryotes than the other giant viruses and has been recently classified as an independent species. However, details of its morphology and maturation process in host cells remain unclear. Here, we investigated the particle morphology of medusavirus inside and outside infected cells using conventional transmission electron microscopy (C-TEM) and cryo-electron microscopy (cryo-EM). The C-TEM of amoeba infected with the medusavirus showed four types of particles: empty, DNA-full, and the corresponding intermediates. Time-dependent changes in the proportion and following intracellular localization of these particles suggested a new maturation process for the medusavirus. Empty particles and viral DNAs were produced independently in the cytoplasm and nucleus, respectively, and only empty particles located near the nucleus incorporated the viral DNA into the capsid. All four types of particles were also found outside the cells. The cryo-EM of these particles showed that the intact capsid structure, covered with three different types of spikes, was conserved among all particle types, although with minor size-related differences. The internal membrane exhibited a structural array similar to that of the capsid, interacted closely with the capsid, and displayed open membrane structures in the empty and empty-intermediate particles. This result suggests that the open structures in the internal membrane are used for an exchange of scaffold proteins and viral DNA during the maturation process. This new model of the maturation process of medusavirus provides insight into the structural and behavioral diversity of giant viruses.

**Importance:** Giant viruses exhibit diverse morphologies and maturation processes. In the present study, medusavirus showed four types of particle morphologies both inside and outside the infected cells, when propagated in the laboratory using amoeba culture. Time-course analysis of the medusavirus particles in the infected cells reveals a new maturation process. Empty viral particles and viral DNAs were produced independently in the cytoplasm and nucleus, and only the empty particles located near the nucleus incorporated the viral DNA. Consequently, many immature particles, along with mature virions, were released from the host cells. Except for showing a small change in size, the capsid structures were well preserved during the maturation process. The empty viral particles and corresponding intermediates showed open membrane structures, which are presumably used for exchanging scaffold proteins and viral DNAs.

## Introduction

Medusavirus is a giant virus isolated from a hot spring water source in Japan (1), and subsequently propagated in *Acanthamoeba* culture similar to other giant viruses reported to date (2–5). Because of its ability to convert host amoeba cells into a cyst, the virus was named after the mythical monster “Medusa”. Medusavirus has a genome of 381 kb, which encodes 461 putative proteins, 86 of which have their closest homologs in *Acanthamoeba*. Furthermore, the genome of its laboratory host, *Acanthamoeba castellanii*, encodes many medusavirus homologs, including the major capsid protein (MCP), suggesting that amoebae are the most promising natural hosts of medusavirus since ancient times, and lateral gene transfers have repeatedly occurred between the virus and amoeba (1). The genome of medusavirus encodes a complete set of histone proteins: four core histones and one linker histone. Recently, a sister strain of medusavirus, named *Medusavirus stheno*, was isolated from a river near Kyoto, Japan, in which histone H3 and H4-encoding genes are fused together (6). Furthermore, the DNA polymerase of medusavirus is phylogenetically placed at the root of the eukaryotic clade. This suggests that medusavirus is closer to eukaryotes than to other viruses (1).

Medusavirus taxonomically belongs to the phylum Nucleocytoviricota (7), which is traditionally referred to as the nucleocytoplasmic large DNA virus (NCLDV), an expansive group of double-stranded DNA (dsDNA) viruses that possess a large capsid (>190 nm) and genome (100 kb to 2.5 Mb) and infect various eukaryotic hosts (8, 9). Currently, the following families are classified into NCLDVs: *Ascoviridae*, *Asfarviridae*, *Iridoviridae*, *Marseilleviridae*, *Mimiviridae*, *Phycodnaviridae*, *and Poxviridae* (10). Medusavirus was classified as an independent species in NCLDVs.

Cryo-electron microscopy (cryo-EM) single particle analysis (SPA) using a 200-kV electron microscope showed that the medusavirus virion is composed of an icosahedron with triangulation number (11) T = 277 (h = 7, k = 12) and exhibits a diameter of 260 nm between the opposing five-fold vertices (1). The 8-nm single-layered major capsid is covered with 14-nm spherical-headed spikes extending from each capsomere. The viral capsid is backed with a 6-nm thick internal membrane, which encloses the viral DNA, as is commonly found in NCLDVs. The membrane extruded to the five-fold vertices of the icosahedral capsid interacts directly with the capsid. The medusavirus particles purified in the laboratory are either filled with DNA or contain no DNA inside; however, both types of particles are structurally similar, as shown by the ~30-Å resolution cryo-EM maps.

Many NCLDVs form a local compartment within the host cytoplasm, called the viral factory, where the virus propagates efficiently (12). Medusavirus does not form such a viral factory in the host cytoplasm for virus replication (1). Fluorescent in situ hybridization (FISH) analysis of the medusavirus-infected cells revealed that the viral DNA first localized to the host nucleus. Then, the signals of the newly synthesized medusavirus DNA gradually became stronger at the periphery of the nucleus up to 8 h post-infection (hpi), indicating that the replication of the viral DNA starts in the host cell nucleus. At 14 hpi, the viral DNA signal spread to the cytoplasm of the host cells. The newly born virions were then released from host cells. A recent study suggests that early expression of genes including histone H1 of medusavirus results in the remodeling of the host nuclear environment before medusavirus DNA replication (13). Together, these observations suggest that medusavirus exhibits unique characteristics with respect to particle maturation compared with other NCLDVs. However, the morphological features of the medusavirus during maturation have not yet been investigated.

In the current study, we found four types of medusavirus particles (DNA-empty, DNA-full, and their corresponding intermediates) inside and outside the virus-infected amoeba cells. Time-course analysis of medusavirus-infected cells using conventional transmission electron microscopy (C-TEM) suggested that these four types of particles show the maturation process in the cell. Furthermore, the intracellular localizations of different types of medusavirus particles indicated that the empty particles and the viral DNAs are produced independently in the cytoplasm and nucleus, respectively, and only the empty particles located near the nucleus incorporate viral DNA into the capsid. Consequently, many empty particles, along with DNA-full particles, are released from the host cell. Cryo-EM using a 300-kV electron microscope revealed that the medusavirus capsid was covered with three types of uniformly arranged spikes and the capsid structure was well conserved throughout the maturation process, except for a small difference in size, depending on the presence or absence of DNA. The internal membrane exhibited a structural array pattern similar to that of the capsid and included an open membrane structure in the empty particles. The open structures in the internal membrane are presumably used for exchanging scaffold proteins and viral DNA during the maturation process. Based on these observations, we propose a new maturation model for medusavirus, which provides insight into the structural and behavioral diversity of giant viruses.

## Results

### Formation of four types of medusavirus particles in the host cell during maturation

Amoeba cells infected with medusavirus were chemically fixed, embedded in plastic resin, and sliced into thin sections, which were then examined by C-TEM. Many replicated medusavirus particles were observed in the host cell cytoplasm at 22 hpi (Fig. 1A). The particles were classified into four different types, depending on their structure: empty (i.e., lacking DNA: Empty), DNA-full (Full), Empty-intermediate (E-I), and Full-intermediate (F-I) (Fig. 1B). The E-I particles were filled with a low-density material (not viral DNA), whereas the Empty particles exhibited an empty capsid. The F-I particles were partially filled with viral DNA, whereas the Full particles were completely filled with viral DNA. These particles were distributed in the cytoplasm at a certain rate, depending on the time point post-infection.

**Figure 1.**
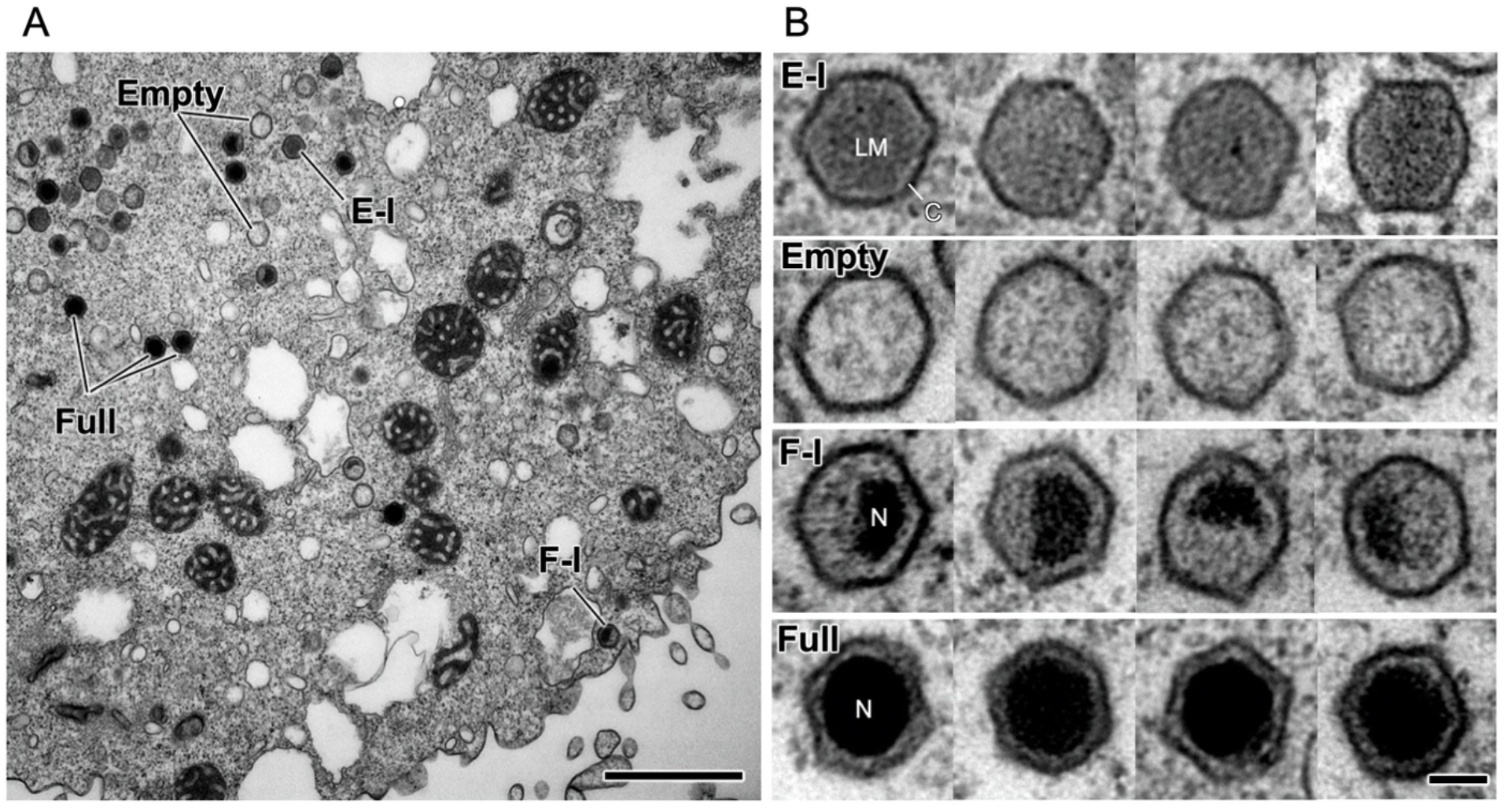
C-TEM image of amoeba cells infected with medusavirus. A) A representative micrograph at 22 h post-infection (hpi). Empty (E), Empty-intermediate (E-I), Full (F), and Full-intermediate (F-I) particles are labeled. B) Zoom-in images of the four different types of medusavirus particles. E-I, capsid is filled with a low-density material; Empty, empty capsid; F-I, partially filled with viral DNA; Full, capsid is completely filled with viral DNA. LM indicates a low-density material. C indicates a viral capsid. N indicates a viral nucleoid. Scale bars = 1 μm in A, and 100 nm in B.

To determine the progression of one type of medusavirus particle to another during the maturation process, the medusavirus-infected amoeba cells were fixed every 2 h, and thin sections were observed by C-TEM. Figure 2 shows the change in the relative proportion of the four types of newly replicated medusavirus particles in the host cells over time. Data from 0 to 8 hpi were omitted because sufficient viral particles were not observed in the cytoplasm. The Empty and E-I particles (blue and green, respectively, in Fig. 2) accounted for more than 85% of all medusavirus particles up to 14 hpi; their proportion decreased significantly to less than 72% at 16 hpi and then increased again to more than 80% after 18 hpi. This oscillation suggested that the first major duplication and release of medusavirus virions occurred at 16 hpi. This result is consistent with a previous study, in which FISH analysis revealed that the newly synthesized viral DNAs are transferred to the cytoplasm at 14 hpi, and then virions appear extracellularly (1). The temporary increase in the number of Empty particles and decrease in the number of E-I particles at 14 hpi were probably the result of the conversion of E-I particles into Empty particles by the release of low-density material. The proportion of F-I and Full particles (red and yellow, respectively, in Fig. 2) increased drastically at 16 hpi, whereas that of Empty particles decreased. This showed that the viral DNA was packaged in the empty capsid at this stage. Mature virions were released from host cells at 16 hpi. After 18 hpi, the proportions of all four types of medusavirus particles gradually reached a constant level, suggesting that the replication process of the late-infecting viruses overlaps with that of the early-infecting viruses. Together, these observations suggest that the medusavirus maturation process proceeds as follows: (1) formation of E-I particles is carried out in the host cytoplasm; (2) conversion of E-I particles into Empty particles by the release of low-density material from the capsid; (3) incorporation of viral DNA produced in the host nucleus into the capsid of Empty particles; and (4) production of Full particles *via* the F-I stage. The temporary decrease in the proportion of E-I particles at 14 hpi, followed by a rapid recovery at 16 hpi, indicates a continuous production of procapsids (E-I) in the cytoplasm.

**Figure 2.**
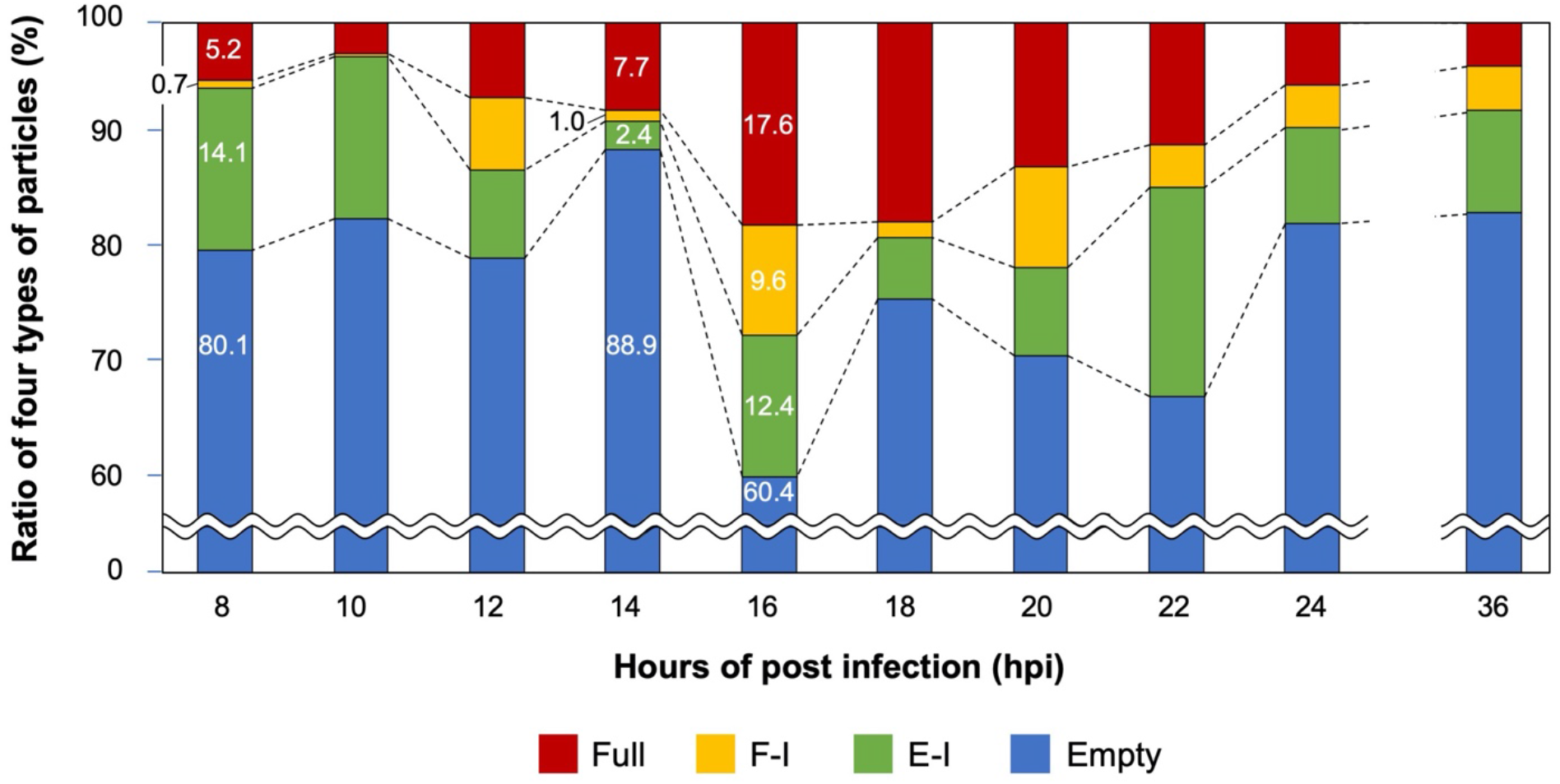
Time-dependent changes in the four types of replicated medusavirus particles in the infected host cells. Proportions of Full, F-I, E-I, and Empty particles are indicated in red, yellow, green, and blue, respectively. The data before 8 hpi and from 24 to 36 hpi were omitted, as the number of particles was insufficient or the morphological changes in the four types of medusavirus particles were not significant.

Next, to determine where the empty medusavirus capsids incorporate viral DNA, we investigated the intracellular localization of Empty and Full particles. The distance of individual medusavirus particles from the host nuclear surface was measured, and the distribution of Empty and Full medusavirus particles was displayed on a chart (Fig. 3). To avoid geometrical interference with the plasma membrane, only particles located at least 2 μm away from the plasma membrane are plotted on the chart. Assuming that the viral particles were randomly distributed in the cytoplasm (random particle distribution), the number of particles increased linearly with the increase in distance from the nuclear surface (dotted line in Fig. 3). The increase in Empty particles was significantly consistent with the random particle distribution curve (white bars in Fig. 3), indicating that the Empty particles are randomly located in the cytoplasm. On the other hand, Full particles were predominantly located at a distance of 0.5-1.0 μm from the nuclear surface, suggesting that Full particles (virions) were generated near the host nucleus surface by incorporating viral DNA in this region.

**Figure 3.**
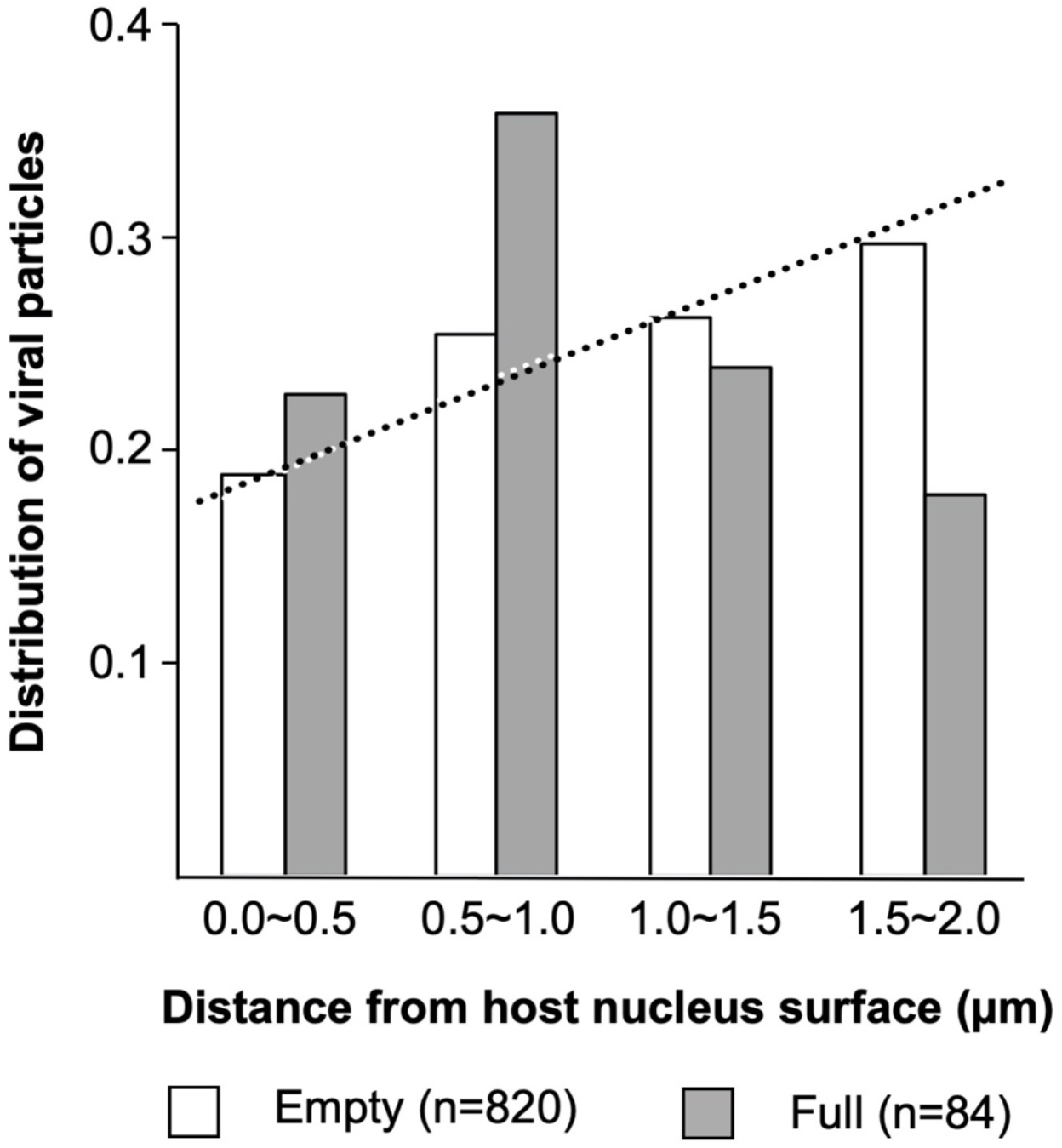
Intracellular distribution of Empty and Full medusavirus particles. The distribution of Empty and Full medusavirus particles is shown with respect to their distances from the host nucleus. The dotted curve shows the distribution of medusavirus particles assuming that they were randomly distributed in the host cell cytoplasm.

### Cryo-EM observations of medusavirus particles outside the host cells

Medusavirus particles were collected from the supernatant of medusavirus-infected amoeba cultures and observed by Cryo-EM. The results clearly showed four types of medusavirus particles (Fig. 4A, Fig. S1A), similar to those identified inside the virus-infected host cells (Fig. 1B). Cryo-EM imaging, which did not involve chemical fixation and dehydration, revealed not only the fine capsid structure but also the morphological differences among the four types of the medusavirus particles (Fig. 4A, Fig. S1A). The E-I particles were filled with low-density material, which was absent from the Empty particles. Interestingly, the internal membrane of E-I and Empty particles was discontinuous (dotted yellow lines in Fig. 4A), whereas that of F-I particles (with the partly incorporated DNA) was deformed (yellow arrows in Fig. 4A). The relative proportions of E-I, Empty, F-I, and Full particles outside the host cells were 28%, 22%, 8%, and 38%, respectively (Fig. 4B); these fractions were significantly different from those inside the host cells (Fig. 2). The Empty and E-I particles (blue and green, respectively, in Fig. 4B) together represented 50% of all medusavirus particles outside the host cells, which is considerably lower than their proportion inside the host cells (>80%), except at 16 hpi. By contrast, Full and F-I particles (red and yellow, respectively, in Fig. 4B) constituted 46% of all medusavirus particles outside the host cells and less than 20% of all medusavirus particles inside the host cells, except at 16 hpi. The fraction of mature particles (virions) outside the host cells was higher than that inside of the cells, suggesting that virions are selectively released from the cells.

**Figure 4.**
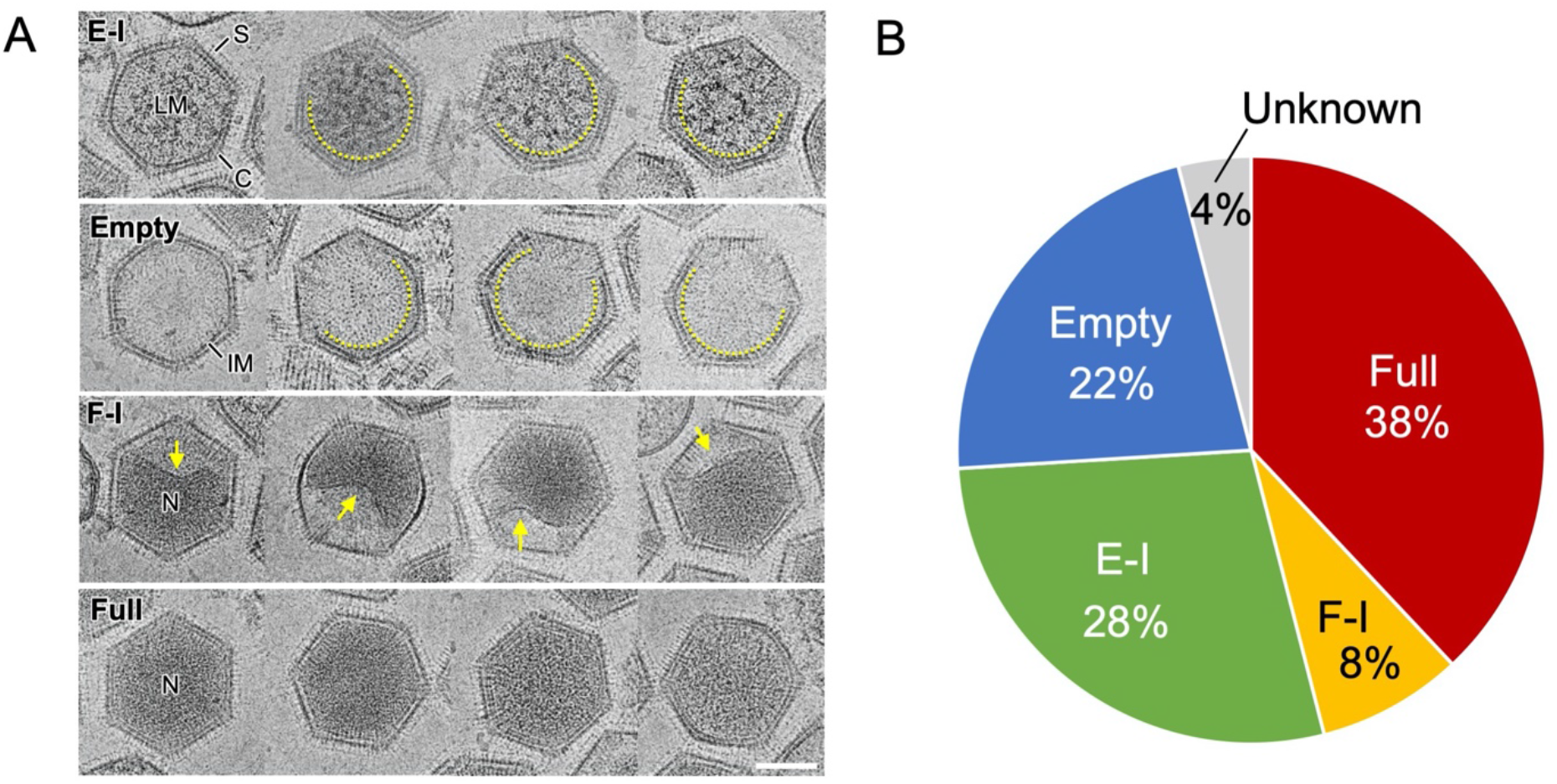
Four types of medusavirus particles observed outside the host cells. A) Cryo-EM images of four different types of medusavirus particles: E-I, Empty, F-I, and Full. Internal membrane (IM) showed a discontinuous structure in the E-I and Empty particles (dotted yellow curves). S: spike, C: capsid, LM: low-density material, N: nucleoid. Scale bar = 100 nm. B) Fractions of the four types of medusavirus particles outside the host cells. The “Unknown” category represents the unclassified broken particles.

### Visualization of open structures in the internal membrane by cryo-electron tomography

The discontinuous internal membrane (dotted yellow curves in Fig. 4A) is reminiscent of the formation of open structures in the Empty and E-I particles. Cryo-electron tomography (cryo-ET) was used to investigate the three-dimensional (3D) structure of the discontinuous internal membranes in E-I and Empty medusavirus particles (Fig. 5, Movies S1 and S2). The internal membrane was discontinued at one or two places in Empty particles (arrowheads in Fig. 5A and B) but was completely closed in Full particles. A representative Empty particle was selected and segmented in the tomogram (Fig. 5B and D). The results of cryo-ET showed that the Empty particle contained an open structure, whereas the external capsid was closed. It has been suggested that the open structure of the internal membrane in Empty particles is used for exchanging scaffold proteins and viral DNA, although how these molecules pass through the external capsid is not clear at this resolution. On the other hand, cryo-ET of a representative F-I particle (Fig. 5C and E) showed that the internal membrane was partly detached from the external capsid and deformed, but it continuously surrounded the viral DNA. However, in F-I particles, how the remaining viral DNA is acquired through the closed internal membrane remains unclear.

**Figure 5.**
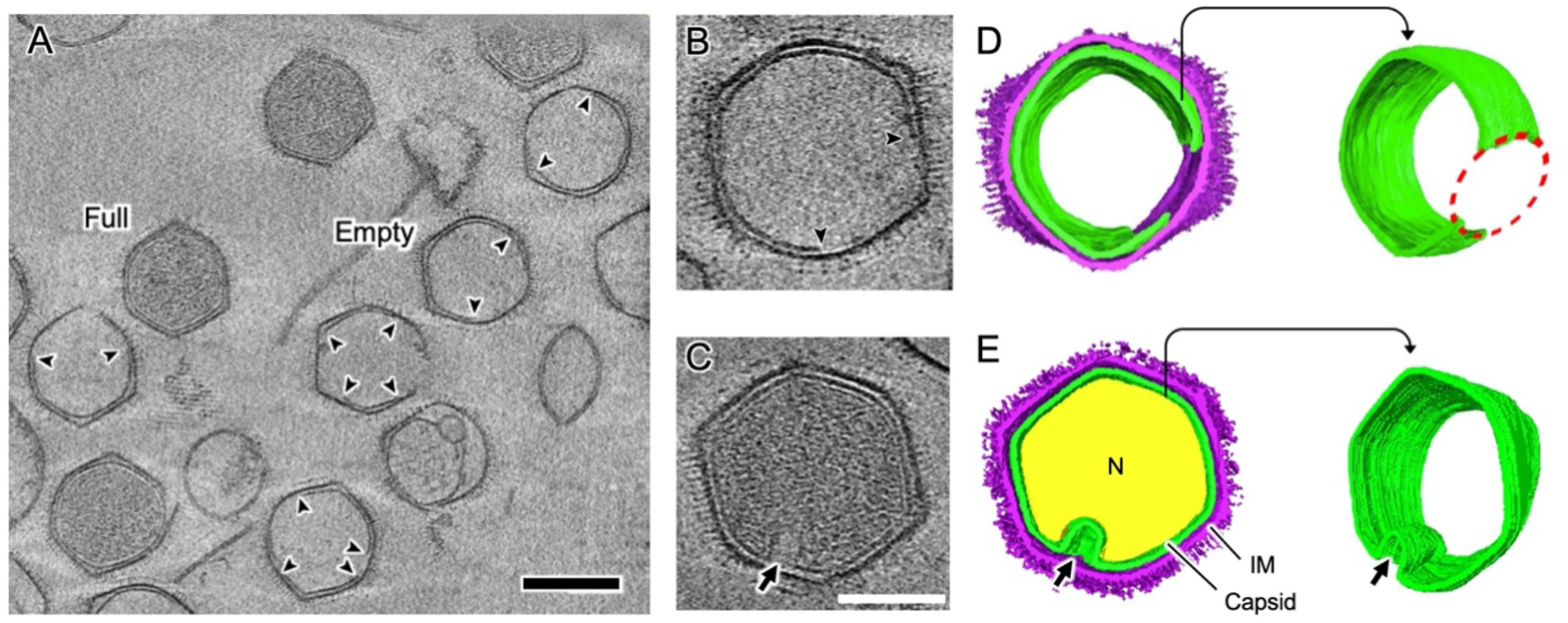
Structural analysis of the internal membrane of medusavirus particles by cryo-electron tomography. A) A tomogram slice of the different types of medusavirus particles outside the host cells. The internal membranes of the Empty and E-I particles were discontinuous (arrowheads). The representative Full and Empty medusavirus particles are labeled. Scale bar = 200 nm. B, D) A tomogram slice (B) and its segmented volumes (D) of the E-I medusavirus particle. The discontinuous internal membrane shows an open structure in the membrane (red dotted circle). C, E) A tomogram slice (C) and segmented volumes (E) of the F-I medusavirus particle. The internal membrane is partially detached from the external capsid and deformed (arrow), but the membrane is completely closed. Capsid, internal membrane (IM), and nucleoid (N) are indicated in purple, green, and yellow, respectively. Scale bars = 200 nm in A, 100 nm in C.

### Cryo-EM SPA of Empty and Full particles

The cryo-EM SPA of Empty and Full medusavirus particles was performed using a 300-kV microscope by imposing icosahedral symmetry. A total of 2,000 micrographs were analyzed (Fig. S1A), and 4,551 Empty particles and 6,981 Full particles were manually selected for 3D reconstruction (Table 1). The two-dimensional (2D) class averages clearly showed whether the capsid of Empty and Full particles was empty or contained viral DNA (Fig. S1B and C). The resolution of these 3D maps was finally estimated at 21.5 Å for Empty particles and 19.5 Å for Full particles using the gold standard Fourier Shell Correlation (FSC) (Fig. S1D and E). As shown in the tomography map (Fig. 5), the medusavirus particles were relatively flexible, limiting the image resolution that could be achieved for both the Empty and Full particles. However, the cryo-EM maps provided detailed structural information for the medusavirus particles as described below.

**Table 1.** Cryo-EM dataset

|  |  |  |
| --- | --- | --- |
| Microscope | Titan Krios G3 |  |
| Accelerating voltage (kV) | 300 |  |
| Spherical aberration (mm) | 0.1 (Cs corrector) |  |
| Detector | Falcon III |  |
| Total dose (e<sup>-</sup>/Å<sup>2</sup>) | 30 |  |
| Micrographs | 2,084 |  |
| Frames per micrograph | 40 |  |
| Nominal magnification | 22,500× |  |
| Pixel spacing on the specimen (Å/pixel) | 3.03 |  |
|  | Empty particle | Full particle |
| Initial particles picked | 4,625 | 7,038 |
| Final particles used | 4,551 | 6,981 |
| Symmetry imposed | I1 | I1 |
| Resolution (Å) | 21.5 | 19.5 |

The first prominent structural feature was the presence of long or wide spikes that extend around the vertices of the capsid surface (asterisks and arrows in Fig. 6A–D), in addition to the regular spikes that covered the entire capsid. The long spikes were 27 nm in length (measured from the top of the MCP trimer) and 6 nm in width (same as the regular spikes) (magenta in Fig. S2A and B). On the other hand, the width of wide spikes was 9 nm at the root, and their length was 13 nm (same as the regular spikes) (green in Fig. S2A and B). Interestingly, these special spikes were located separately between the pentasymmetron and trisymmetron (Fig. S2A and C). The triangular asymmetrical unit of the pentasymmetron consisted of six capsomers (MCP trimer) labeled P_1_–P_6_, and two wide spikes located in P_3_ and P_6_ (Fig. S2A and C). The third wide spike (TW in Fig. S2A and C) was located in the trisymmetron adjacent to the P_6_ wide spike, whereas the long spike (TL in Fig. S2A and C) was located in the trisymmetron adjacent to the P_5_ regular spike. The wide spikes showed a wider root underneath the spherical head (green in Fig. S2A and B). The long spikes showed an extended tail underneath the spherical head, although the density was weak (asterisks in Fig. 6A and B, asterisk in Fig. S2B). The spherical heads of both long and wide spikes were wider than those of regular spikes (Fig. S2B). All types of spikes were found in both Empty and Full medusavirus particles (Fig. 6A–D), suggesting that spikes are formed at the early maturation stages. These spikes are possibly involved in host recognition and host membrane permeation, although further investigation is needed to verify their function.

**Figure 6.**
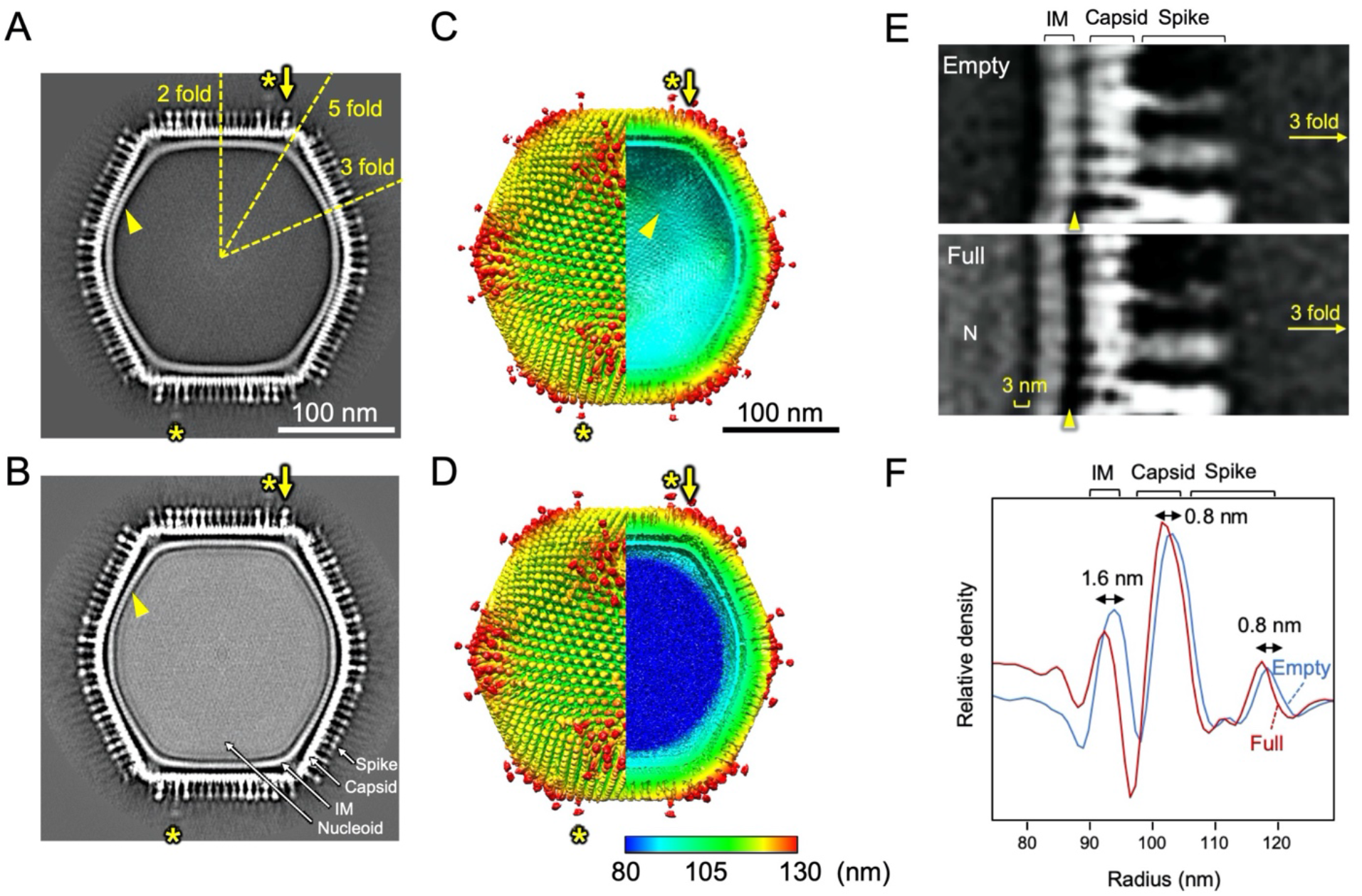
Cryo-EM 3D reconstructions of Empty and Full particles and their size comparison. A-D) Center slices and surface renderings of the cryo-EM maps of the Empty (A, C) and Full (B, D) particles. The 3D maps were colored by radius. Asterisks indicate long spikes. Arrows indicate wide spikes. E) Center slices of the Empty (top) and Full (bottom) particle cryo-EM maps near the three-fold axis. Gaps between the capsid and internal membrane and between the internal membrane and nucleoid are indicated by yellow bracket and yellow arrowhead, respectively. F) Radial profile of the Empty (blue) and Full (red) particles. The size differences in the internal membrane (IM), capsid, and spikes are indicated.

The second prominent structural feature was the lattice array on the internal membrane of the medusavirus particles (arrowhead in Fig. 6A and C), which showed similarity to the array of the external viral capsid (Fig. S3). The internal membrane had a width of 5.7 nm (Fig. S4A), which is typical of lipid bilayers, and looked similar to the internal membrane of other NCLDVs (14–16). However, the lattice array on the internal membrane of medusavirus was unique and structurally reminiscent of the inner protein core shell of faustovirus (17) and ASFV (Fig. S5B) (15, 16). To ensure that the internal membrane consists of a lipid membrane rather than a protein shell, medusavirus particles were treated with 50% ethyl alcohol. The 2D averaged images of the Empty medusavirus particles showed that the internal membrane disappeared following the alcohol treatment (Fig. S4B), suggesting that the internal membrane is composed of a lipid membrane rather than a protein shell. Based on these results, we concluded that the internal membrane of medusavirus consists of a lipid bilayer containing a membrane protein, rather than only lipid, and exhibited a structural array pattern (IM in Fig. S5). The membrane proteins potentially allow the formation of open structures, as observed by cryo-ET, in the internal membrane (Fig. 5). Furthermore, a gap of approximately 3 nm was observed between the internal membrane and the nucleoid (yellow bracket in Fig. 6E), which has not been reported in other NCLDVs. This gap may be filled with low-density materials, and the internal membrane indirectly encloses the viral nucleoid.

Unexpectedly, the Empty and Full medusavirus particles showed highly similar capsid structures but slightly different particle sizes (Fig. 6F). Compared with Empty particles, the Full particles showed a 0.8 nm reduction in particle radius at the spike edges and a maximum radius reduction of up to 1.6 nm at the internal membrane region. The cryo-EM maps clearly showed that the gap between the internal membrane and the capsid was greater in the Full particles than Empty particles (arrowheads in Fig. 6E). This suggests that the strong interaction between the incorporated viral DNA and the internal membrane caused virus particle contraction.

## Discussion

In the present study, we first investigated the maturation process of medusavirus by performing a time-course analysis of medusavirus-infected amoeba cells with C-TEM. We found four different types of particles inside the cells, namely, E-I, Empty, F-I, and Full particles, which demonstrate the medusavirus maturation process in this sequence inside the host cell. These particles were also found in the supernatant of the amoeba culture, and the fine structures of Empty and Full particles were examined and compared by cryo-EM SPA, showing that the capsid structure is well preserved during the maturation process, except for a small change in size. Overall, this study reveals a new particle formation and maturation process of medusavirus.

The capsid of E-I medusavirus particles was filled with a low-density material rather than viral DNA at the early stage of maturation (Fig. 2). The low-density material is reminiscent of a scaffold protein initially reported in dsDNA phages during capsid assembly (18). These phages use scaffold proteins to build a uniform icosahedral capsid and then release the proteins after the assembly of the capsid is complete and before the viral DNA is incorporated into the capsid. In NCLDVs such as mimivirus (19), ASFV (20), vaccinia virus (21), and PBCV-1 (22), the formation of virus particles begins with a membrane sheet scaffolded with proteins, and the resultant curled membrane sheet is used as a template for capsid assembly. Viral DNAs are then incorporated into the capsid by replacing the scaffold protein before the capsid is completely closed. All these processes occur at the periphery of the viral factory. In the case of medusavirus, which does not form a viral factory, such a capsid formation process has not yet been observed in the host cytoplasm, but a capsid assembly process similar to that of NCLDVs is believed to occur in medusavirus using scaffold proteins. The major difference compared with other NCLDVs is that the medusavirus produces many empty capsids inside and outside the host cell. The open structure in the internal membrane of the Empty particles (Figs. 4 and 5) may allow the release of scaffold protein and the uptake of viral DNA, although the underlying mechanism remains unclear.

Careful observation of the open membrane structure in the medusavirus capsid revealed that the membrane opening always occurred near the outer capsid (Figs. 4A and 5). The cryo-EM map of medusavirus showed an array pattern on the internal membrane similar to that on the outer capsid (Figs. 6 and S3), suggesting that membrane proteins contained in the internal membrane strongly interact with the outer capsid. PBCV-1 and ASFV, whose structures were reported at a higher resolution (14, 15), exhibited a smooth internal membrane (Fig. S5) and did not show an open membrane structure. Membrane proteins in the internal membrane are likely to control the open structures in medusavirus by collaboratively interacting with the outer capsid.

The cryo-EM reconstructions of the medusavirus at ~20 Å resolution revealed no significant structural differences between the Empty and Full particles other than the presence or absence of the viral DNA, but the sizes of the two particles showed a slight difference (Fig. 6E and F). Interestingly, the average radius of the internal membrane was less in Full particles (by 1.6 nm) than in Empty particles, whereas the radii of the capsid and outermost spikes were shorter in

Empty particles (by 0.8 nm) than in Full particles. This suggests that the internal membrane strongly interacts with the nucleoid, causing a major contraction in the internal membrane region. However, the space between the internal membrane and the nucleoid was separated at a distance of 3 nm (yellow bracket in Fig. 6E), and there are no interconnecting components. On the other hand, a minor contraction of the radii of the capsid and the outermost spikes (by 0.8 nm) also suggests interactions between the internal membrane and outer capsid. The cryo-EM maps showed several interconnecting components between these two structures (Fig. S6). At the five-fold vertices in medusavirus particles, several thread-like structures were observed between the extruded internal membrane and the capsid in both Empty and Full particles (arrows in Fig. S6A and B). Around the two-fold axes of the medusavirus particle surface, in addition to the thread-like structures (arrows in Fig. S6E and F), two symmetrically related two interconnecting structures were clearly identified in both Empty and Full particles (arrowheads in Fig. S6C-F). These interconnecting components form the relatively flexible interactions between the internal membrane and the external capsid.

The cryo-EM SPA of medusavirus particles revealed for the first time long and wide spikes, in addition to the normal spikes. The asymmetric unit showed three wide spikes: two on the pentasymmetron and one on the adjacent trisymmetron (green in Fig. S2). On the other hand, one long spike in the asymmetric unit was adjacent to the wide spike on the trisymmetron (magenta in Fig. S2). These different types of spikes were located adjacent to each other, but no interaction was found between these spikes. Xiao and coworkers reported that in CroV and PBCV-1, the orientation of the MCP trimers is constant in each trisymmetron and in each asymmetric unit of pentasymmetron, except for the MCP trimers located at five corners of the pentasymmetron, and these MCP trimer arrays are interconnected in 60°orientation, eventually forming a large protein cage (23). This characteristic MCP trimer array has also been confirmed in ASFV (15) and Marseillevirus (24), and our results showed that medusavirus also employs this scheme of capsomere arrays (Fig. S2C). The resultant MCP trimer arrangement suggests that the wide spikes in the asymmetric unit of the pentasymmetron are oriented at 60° from the wide and long spikes in the adjacent trisymmetron. The function of these special spikes is currently unknown, and further investigation is needed to understand the biological significance of the spike orientation and function.

The intracellular localization of Full particles was predominantly high near the host nucleus (Fig. 3), suggesting that the DNA packaging of medusavirus occurs in the region surrounding the host nucleus. Medusavirus does not form a viral factory in amoeba (1). These results suggested that empty particles and viral DNA are independently produced in the cytoplasm and nucleus, respectively, of the host cells and the Empty particles incorporate the viral DNA near the host nucleus. Eventually, DNA packaging is relatively inefficient in medusavirus compared with other NCLDVs that form a viral factory (12). In iridovirus (25) and PBCV-1 (22), which form a viral factory near the host nucleus, capsid production occurs in the viral factory, and viral DNA replicates inside the host nucleus and is delivered to the viral factory for encapsulation by the capsid. Mimivirus (19) and vaccinia virus (26), which also form a viral factory, produce the capsids around the viral factory and incorporate the viral DNA which is replicated in the viral factory itself. Thus, because of the viral factory, these NCLDVs package DNA with greater efficiency than medusavirus, in which DNA packaging accidentally occurs near the host nucleus because of the random distribution of Empty particles in the cytoplasm (Fig. 3). Consequently, the medusavirus host releases many immature particles, including E-I, Empty, and F-I particles, from the host cells in addition to the matured Full particles (Fig. 4). However, the ratio of Full particles outside the host cells (38%) is significantly higher than that in the host cell cytoplasm (<17.6%) (Figs. 2 and 4B). Medusavirus may have a mechanism for selectively releasing the mature virions. Overall, these results indicate that medusavirus employs a relatively primitive strategy for maturation, although its genome is phylogenetically closer to eukaryotes than the other giant viruses (1).

Based on our observations of medusavirus, we propose a new maturation process for giant viruses (Fig. 7): (1) The E-I particles filled with low-density material are initially produced in the cytoplasm using the host intracellular membranes; (2) the E-I particles then turn into Empty particles by releasing the low-density material (scaffold protein); (3) the Empty particles located near the host nucleus incorporate viral DNA produced in the host nucleus independently, thus converting into F-I particles; (4) F-I particles encapsulate all of the viral DNA to form Full particles. The open structures of the internal membrane are likely used to exchange the scaffold proteins with the viral DNA, but the underlying mechanism remains unknown. (5) The open structures in the internal membrane can be closed with an extra membrane brought into the capsid along with the viral DNA. (6) Finally, the mature Full particles, together with immature particles of E-I, Empty and F-I particles, are released from the amoeba cell. The cryo-EM SPA of medusavirus particles outside the host cells showed that all four types of particles possess a perfect set of spikes on the capsid. In general, the spikes of progeny viruses are inactive inside the host cell to avoid a reaction with the host and must be activated before the release from host cells, as reported for Dengue and Zika viruses (27, 28). Small structural modifications in the spikes may occur at the molecular level, or the spike structures inside the host cells may be different from those outside the host cells; however, these structural modifications cannot be identified at the currently achievable resolution and with the current methodology. Increase in the proportion of Full and F-I medusavirus particles outside the host cells may be a result of a structural modification in the spikes, which triggers the selective release of mature particles. Further investigation is necessary to clarify the mechanisms underlying the medusavirus maturation process.

**Figure 7.**
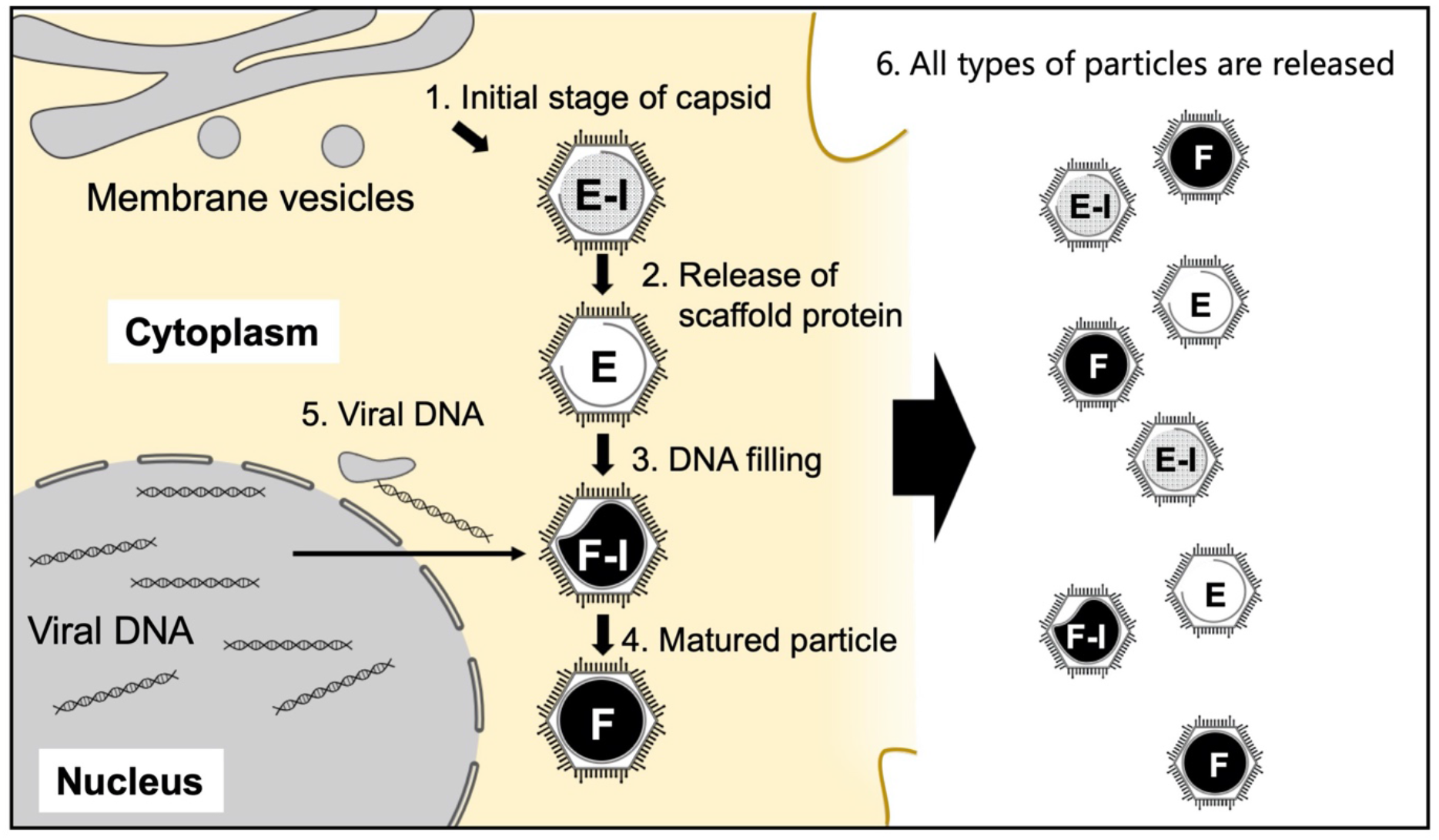
Proposed maturation process of medusavirus. (1) E-I particles are initially generated in the cytoplasm. (2) E-I particles change to Empty particles by releasing the low-density materials (scaffold proteins). (3) Empty particles incorporate viral DNA near the host nucleus (F-I and Full particles). (4) Full particles are generated. (5) The open structures in the internal membrane can be closed with an extra membrane brought into the capsid along with the viral DNA. (6) Mature visions are released from the host cells together with the immature E-I, Empty and F-I particles.

In our first report, the Full and Empty medusavirus particles were described at 31.3 Å and 31.7 Å resolution, respectively, by cryo-EM SPA using a 200-kV microscope (1). The cryo-EM maps determined the triangulation number of the medusavirus icosahedral capsid as 277 and revealed the spiky feature of the capsid and the characteristic extrusion of the internal membrane structure into the five-fold vertices of the capsid. In the current study, the resolution was improved to 19.5 Å and 21.5 Å in Full and Empty particles, respectively, with a 300-kV microscope. The new cryo-EM maps revealed the presence of long and wide spikes in addition to the regular spikes and a membrane protein array in the internal membrane and its connection with the outer capsid. The sub-nanometer structures of the icosahedral giant viruses have been reported in recent years, including the 3.5-Å structure of PBCV-1 (14), 4.6-Å and 4.8-Å structures of ASFV (15, 16), 4.4-Å structure of Melbournevirus (24), 7.7-Å structure of tokyovirus (29), and 8.6-Å structure of Singapore grouper iridovirus (30). These cryo-EM single particle reconstructions were generated using 300 or higher kV microscopes to avoid the defocus gradient and to keep enough electron transparency for large objects (31). Furthermore, some image processing techniques such as Ewald sphere correction (32), magnification anisotropy, and higher order aberration correction (33) were applied to improve the resolution. For structures with resolution exceeding 5 Å, a method of block-based reconstruction was also adopted (34). In these cryo-EM SPAs, the initial map resolution, however, reached approximately 1 nm without the image correction techniques. By contrast, the initial resolution of the cryo-EM maps of medusavirus is much lower than 1 nm, mainly because of the distortion of the capsid. Compared with giant viruses with higher resolution structures, the capsid structures of medusavirus with spikes are relatively flexible, degrading the achievable resolution. To obtain flexible domain structures of proteins, a “focused classification and refinement” image processing technique has been developed and tested (35). This technique may enhance the image resolution of medusavirus in the future.

## Materials and Methods

### Growth and purification of medusavirus

*A. castellanii* strain Neff was used as the viral host. Amoeba cells were cultured at 26°C in flasks containing 100 ml of peptone-yeast-glucose (PYG) medium as described previously (1). A total of four cultures were grown and inoculated with medusavirus at a multiplicity of infection (MOI) of 1–2. The newly born medusaviruses were harvested 3 days post-infection (dpi). Amoeba cells and cell debris were removed by centrifugation (800×g, 5 min, 24°C), and the medusavirus particles were concentrated by an additional centrifugation (8,000×g, 35 min, 4°C). The viral particles were suspended in 10 ml phosphate-buffered saline (PBS) and filtered using a 0.45-μm filter (Millex-AA, Merck Millipore, Darmstadt, Germany). The filtered viral particles were centrifuged (8,500×g, 35 min, 4°C) and resuspended in 10 μl of PBS. This process was repeated 5–10 times to obtain a sufficient amount of medusavirus particles.

### Time-course analysis of infected amoeba cells by C-TEM

Medusavirus-infected amoeba cells were harvested at 2-h intervals and fixed with 2% glutaraldehyde in PBS at a room temperature of 23°C for 30 min. The cells were washed three times with PBS and then post-fixed with 2% osmium tetroxide in PBS at a room temperature of 23°C for 1 h. The fixed cells were dehydrated using an ethanol gradient (50%, 70%, 80%, 90%, 95%, 100% for 5 minutes each) at a room temperature of 23°C. The dehydrated cells were infiltrated with propylene oxide and embedded in epoxy resin mixture (Quetol 812; Nissin EM Co. Ltd., Tokyo, Japan). The resin was polymerized at 60°C for 2 days. Ultrathin sections (approximately 70-nm thickness) were prepared with a diamond knife using an ultramicrotome (EM-UC7; Leica Microsystems, Austria). The sections were stained with 2% uranyl acetate for 5 min and then with 0.4% lead citrate for 1 min. The stained sections were visualized using a JEM1010 microscope (JEOL Ltd., Japan) at an acceleration voltage of 80 kV.

### Cryo-EM and SPA

A 2.5-ml suspension of the purified medusavirus particles was applied to a quantifoil grid (R1.2/1.3 Mo) (Quantifoil Micro Tools GmbH, Germany), which was glow-discharged beforehand for 30 s. The grid was then blotted with a filter paper (blotting time: 7 s, blotting force: 10) and plunge-frozen at 4°C under 95% humidity using Vitrobot Mark IV (Thermo Fisher Scientific, USA). The frozen grid was imaged using a Titan Krios G3 microscope at an acceleration voltage of 300 kV (Thermo Fisher Scientific, USA). Movies were recorded on a Falcon III detector at a nominal magnification of 22,500×, corresponding to 3.03 Å per pixel on the specimen. A low-dose method (exposure at 10 electrons per Å^2^ per second) was used, and the total number of electrons accumulated on the specimen was ~30 electrons per Å^2^ for a 3-s exposure time. A GIF-quantum energy filter (Gatan Inc., USA) was used with a slit width of 20 eV to remove inelastically scattered electrons. Individual micrograph movies were subjected to per-frame drift correction by MotionCor2 (36), and the contrast transfer function parameters were estimated by CTFFIND4 (37).

To perform SPA, 4,625 Empty particles and 7,038 Full particles were manually selected and extracted from 2,084 motion-corrected images using the RELION3.0 software (38). Subsequently, 4,551 Empty particles and 6,981 Full particles were selected from the extracted particles by 2D classification (Fig. S1A and C) and used for 3D reconstruction with imposing icosahedral symmetry. The resolution of the final 3D maps was estimated at 21.5 Å for Empty particle and 19.5 Å for Full particle using the 0.143 gold standard FSC criterion (39) (Fig. S1B and D). The 3D reconstructions were visualized using the UCSF Chimera software (40).

### Radial profile of Full and Empty particles

The five-fold axis of Full and Empty particles was reoriented to the front, and a center slice of the z-axis was extracted from the reoriented volumes. The sliced image was rotationally averaged using the EMAN2 software (41). Then, the radial profile was calculated using the Fiji image analysis software suite (42).

### Cryo-electron tomography

Colloidal gold particles (15 nm) were mixed with the sample as a fiducial marker before freezing. The frozen grid was examined with a CRYO ARM 300 electron microscope (JEOL Inc., Japan) at 300 kV accelerating voltage. Tilt series images were collected in the range from −60° to +60° with a 2° increment step using a low-dose mode, where the total electron dose of 61 images was less than 100 e^-^/Å^2^ on the specimen. Images were recorded on a K3 camera (Gatan Inc., USA) at a nominal magnification of 15,000× and a pixel size of 3.257 Å on the specimen using the batch tomography procedure of the Serial EM Software (43). Image alignment and tomographic reconstruction were performed with the IMOD software version 4.7.15 (44) using fiducial markers. The final tomograms were calculated with the simultaneous iterative reconstruction technique using images of 6.514 Å per pixel after applying a pixel-binning of two. The image segmentation in the 3D reconstructions was performed with Amira version 5.4.5 (Thermo Fisher Scientific, USA).

## Data availability

The density maps have been deposited in the EMDB with the accession codes EMD-32073 (Full particle), EMD-32072 (Empty particle), and EMD-32071 (tomography).

## Acknowledgments

We thank Sachiko Yamada (NIPS) for image segmentation, Naoyuki Miyazaki (formerly Osaka University) for the data acquisition of single particle analysis, and Naoki Hosoki (JEOL Ltd.) for the data acquisition of cryo-electron tomography, Raymond N. Burton-Smith for critical reading of the manuscript. This study was supported by MEXT/KAKENHI under Grant Numbers JP17H05825 and JP19H04845 to K.M., JSPS/KAKENHI under Grant Number 20H03078 to M.T., BINDS from AMED under Grant Number JP18am0101072 (support number 1162) to K.M., the Joint Research of ExCELLS (20-004) to K.M., and the Cooperative Study Program of the National Institute for Physiological Sciences (20-239) to M.T.

## Author contributions

K.M. conceived the project. M.T. provided the medusavirus sample. M.T. and R.W. collected and processed the C-TEM data. C.S. prepared the cryo-EM sample and collected the cryo-EM data. C.S., Y.K., R.W., and K.M. processed the cryo-EM data. R.W. prepared the figures. R.W. and K.M. wrote the main manuscript text. All authors reviewed the manuscript.

